# Optineurin is involved in regulating macrophage responses during mycobacterial infection

**DOI:** 10.1101/2023.09.11.557134

**Authors:** Gopalakrishna Ramachandran, Chaitanya Veena Yeruva, Ghanshyam Swarup, Tirumalai R. Raghunand

**Author notes:** For correspondence Tirumalai R. Raghunand. **Email information:** Gopalakrishna Ramachandran -, Chaitanya Veena Yeruva -, Ghanshyam Swarup. Tata Institute of Fundamental Research, 36/P, Gopanpally Village, Serilingampally Mandal, Ranga Reddy District, Hyderabad, 500046, India (Gopalakrishna Ramachandran). Sanofi Healthcare India Pvt. Ltd., Athvelly, Medchal, Hyderabad, 501401, India.

## Abstract

Autophagy has emerged as a critical innate immune mechanism for host elimination of intracellular pathogens, however, the role of the autophagy receptor optineurin during mycobacterial infection is not fully understood. To address this lacuna, we infected bone marrow-derived macrophages (BMDMs) derived from Optn^+/+^ and Optn^-/-^ mice with *Mycobacterium smegmatis*, and observed the infection outcome at sequential time points. While low multiplicity of infection (MOI) did not show any significant difference between BMDMs from the two groups, at high MOI Optn^-/-^ mice-derived macrophages showed significantly lower colony forming unit counts, as well as lower cell counts at 12 h and 24 h post-infection. Quantification of cell numbers and nuclear morphologies at various time points post-infection indicated a markedly higher cell death in the optineurin-deficient macrophages. Optineurin-deficient macrophages showed significantly lower levels of the autophagosomal protein LC3-II upon infection, indicating a potential role for optineurin in regulating autophagy during mycobacterial infection. Moreover, when stimulated by bacterial LPS, optineurin deficient macrophages, showed altered levels of the inflammatory cytokine pro-IL-1β. These observations taken together suggest a novel regulatory role for optineurin during mycobacterial infection, with its deficiency leading to an impairment in macrophage responses.

## INTRODUCTION

Optineurin is a multifunctional protein that is involved in vesicle trafficking, signal transduction, cell survival, innate immunity, inflammation, and autophagy [1-4]. It functions by interacting with several proteins such as Myosin VI, Rab8, Transferrin receptor, LC3 (MAP1LC3B), TBK1 (Tank Binding Kinase 1), Transcription Factor IIIA, and Ubiquitin [5-10]. Due to its interaction with various molecules that possess diverse functions, a deficiency or mutation affecting its interaction is likely to lead to alterations in cellular homeostasis. Optineurin is also essential for the maintenance of organelle structure and function [11], and its deficiency in cell lines leads to Golgi fragmentation, impaired vesicle trafficking and cell death [12, 13]. The property of optineurin to bind to LC3 and ubiquitinated molecules, simultaneously linking them to autophagosomal membranes, add to its function as an autophagy receptor [7]. Autophagy is an intracellular catabolic process that assists in the maintenance of homeostasis through lysosomal degradation [14]. Post-translational modifications such as phosphorylation of optineurin by TBK1 regulate the autophagic activity of optineurin [7, 15]. Optineurin-mediated selective autophagy prevents neurodegeneration during herpes virus infection and prevents viral proliferation [16, 17]. Intracellular bacterial pathogens such as *Salmonella enterica* which escape into the cytosol, are degraded by the cellular autophagy machinery, and this process is enhanced by phosphorylation of optineurin at Serine-177 [7, 18]. Optineurin is also known to play an important role in neutrophil recruitment and the associated inflammatory response, events critical to the control of bacterial infection [19].

*Mycobacterium tuberculosis*(*M. tb*) the causative agent of human tuberculosis, is one of the oldest coexisting pathogens of humans that still causes over a million deaths annually [20]. The ability of *M. tb* to establish infection is founded on its ability to evade host defence mechanisms [21, 22]. This bacterium has evolved intricate mechanisms to manipulate the host’s immune responses, which allow it to survive and replicate in the hostile macrophage environment. While the molecular details of the mycobacterium-macrophage chemistry continue to be elucidated, host autophagy seems to have a predominant role in the progress of the infection [23, 24]. *M. tb* enters its human host *via* aerosols containing infectious bacilli, that are internalised by the lung’s resident alveolar macrophages [20, 25]. This initiates a cascade of events leading to the production of cytokines, induction of autophagy and other cellular homeostatic mechanisms, which coordinate to contain the infection [20, 26]. However, the status of host immunity, and virulence of the infecting strain, are critical in determining the outcome of infection [27].

The ability to regulate cell death is critical during any microbial infection. Macrophages are the critical component of the innate immune system that serve as the first line of defence against infections [23, 28]. However, *M. tb* has evolved sophisticated strategies that manipulate phagosome trafficking pathways, disrupting normal host cellular microbicidal activities [26]. Consequently, *M. tb* survives in the phagosome and limits its fusion with the lysosome, eventually leading to the death of macrophages and dissemination of the bacteria [29-31]. The three major consequences of a mycobacterial infection are necrosis, apoptosis, and survival of infected macrophages [22, 32], with the kinetics of macrophage death being an important parameter that influences the outcome of infection. In this study, we have investigated the role of optineurin during mycobacterial infection, using bone marrow-derived macrophages (BMDMs) derived from Optn^+/+^ and Optn^-/-^ mice. Based on enumeration of cell numbers, classification of nuclear morphologies and quantifying markers of cell death post-infection, our data suggest that deficiency of optineurin leads to an impairment in macrophage responses to mycobacterial infection, highlighting its novel cytoprotective role in this context.

## MATERIALS AND METHODS

### Isolation of Bone marrow-derived macrophages

All mouse experiments were approved by the Institutional Animal Ethics Committee. Optineurin knockout mice were generated by replacing the Exon 2 of the optineurin gene with a cassette containing a β-galactosidase reporter and neomycin resistance gene as described previously [33]. Genotyping of these mice was performed as described in [34]). Primary bone marrow-derived macrophages (BMDMs) were isolated from the femur, tibia, and humerus of littermate Optn^+/+^ and Optn^-/-^ mice. These long bones were sliced at both the ends and the bone marrow was flushed using sterile chilled PBS using a 23G 1.5-inch needle attached to a 10 ml syringe. The recovered bone marrow tissue was homogenized and cultured in 100 cm^2^ non-adherent dishes for 7 days in Macrophage culture media (80% DMEM +10% L929 conditioned media +10% FBS). After 7 days, the cells were scraped in chilled PBS using a sterile cell scraper, resuspended in macrophage culture media, counted, and plated in 6 well-plates for infection.

### Infection of BMDMs

All infection experiments were approved by the Institutional Biosafety Committee. *M. smegmatis* mc^2^155 was used for all experiments unless otherwise specified. *M. smegmatis* was grown in Middlebrook 7H9 broth and Middlebrook 7H10 agar (Difco) supplemented with albumin dextrose complex (5 g BSA, 2 g glucose and 0.85 g NaCl/L), 0.5% (v/v) glycerol and 0.05% (v/v) Tween 80, at 37°C in a shaker incubator with 15 μg/ml kanamycin. The infection was carried out by resuspending exponentially growing *M. smegmatis* in DMEM + 10% FBS + 10% L929 conditioned medium without antibiotics, after proper declumping and dilution. An infection time of 4 h was allowed unless otherwise specified. Intracellular bacterial growth was measured by performing colony forming unit (CFU) counts, determined at the designated time points by lysing infected cells with 0.1% Triton X-100 followed by dilution plating on Middlebrook 7H10 agar. Cytokine and autophagy levels were assessed by western blotting, and quantification of cell death was performed using fluorescence microscopy.

### Microscopy

Images of macrophage growth and microscopic assessment of cell death were performed with Ziess Axioimager Z1 fluorescence microscope at 10x, 20x and 40x magnification.

### Western blotting

Western blotting was performed as described previously [34]. The signals were developed using the Vilber Lourmat Chemi Doc-5000 instrument and quantified using ImageJ. The antibodies used were as follows: Optineurin (Abcam, Ab23666), LC3B (Enzo life science, ALX80308), Actin (Millipore, MAB1501), Cleaved caspase 3 (CST, #9664), IL-1β (Santa Cruz - sc-52012), and GAPDH (Millipore MAB374).

### Statistical Analysis

The Unpaired t-test was performed to determine levels of significance. A P-value less than 0.05 was considered significant. GraphPad Prism 5 and Microsoft Excel were used for data analysis and graph preparation followed by Adobe Photoshop for figure assembly.

## RESULTS

### Fate of *Mycobacteria* upon infection of *Optn*^+/+^ & *Optn*^-/-^ deficient macrophages

To investigate the role of optineurin (Optn) in mycobacterial infection, bone marrow-derived macrophages (BMDMs) from Optn^+/+^ and Optn^-/-^ mice were infected with *Mycobacterium smegmatis* (*M. smegmatis*), an avirulent saprophytic mycobacterial species, at low (10:1) and high (50:1) MOI. This organism has been extensively used as a surrogate model to study the physiology and virulence mechanisms of pathogenic mycobacteria including *M. tb* [35]. At low MOI, we observed no significant differences in CFU counts in Optn^+/+^ vs Optn^-/-^ macrophages. At high MOI however, at 24 h post-infection, Optn^-/-^ macrophages showed lower mycobacterial counts than Optn^+/+^ macrophages (Fig. 1). The reduction in CFU counts over time, signifies macrophage killing of *M. smegmatis*, a reflection of its non-pathogenic nature.

**Figure 1:**
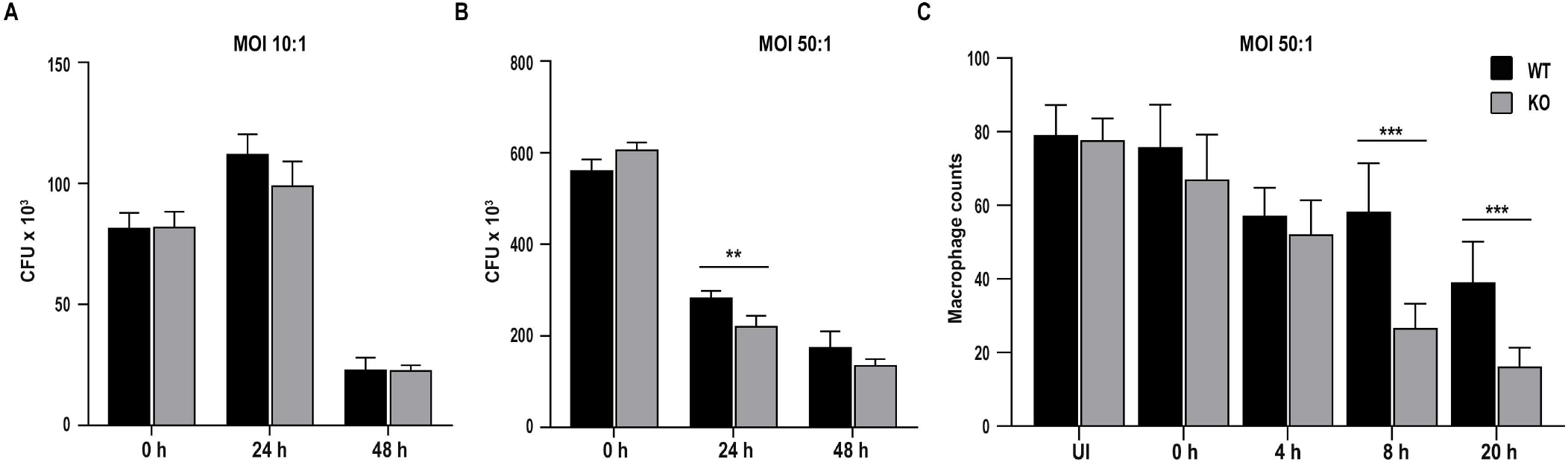
Intracellular viability of *M. smegmatis* in *Optn*^-/-^ macrophages. CFU counts of *M. smegmatis* upon infection of *Optn*^+/+^ and *Optn*^-/-^ bone marrow derived macrophages at an MOI of 10:1 (A) and 50:1 (B). (C) *Optn*^+/+^ and *Optn*^-/-^ BMDM cell counts post 50:1 MOI *M. smegmatis infection* (n=4 independent experiments, minimum 300 cells per experiment per genotype assessed) Bars represent mean ± SD. **p ≤ 0.01, ***p ≤ 0.001.

### *M. smegmatis* induced higher cell death in optineurin-deficient bone marrow-derived macrophages at 50:1 MOI

Based on the above observations, we proceeded to investigate the effect of mycobacterial infection on Optn^-/-^ macrophages using an MOI of 50:1. We found that the loss of cell numbers due to *M. smegmatis* infection was higher in Optn^-/-^ macrophages compared to the Optn^+/+^ macrophages. To quantify this effect, BMDMs were plated onto coverslips, and infected with *M. smegmatis* at an MOI of 50:1. After infection, coverslips with infected macrophages were fixed and stained with DAPI at 4 h,8 h and 20 h time points followed by assessment of total cell count per field. Random fields were selected and cells per field were counted under phase contrast under a 400X magnification. We observed a significantly lower number of Optn^-/-^ macrophages at 8 h and 20 h post infection (Fig. 1C). To gain a deeper insight into the loss of Optn^/-^ macrophages upon infection, we utilized DAPI-stained macrophages for further microscopic examination. On further observation, we noticed alterations in the nuclear morphology of BMDMs post-infection, which were classified into three distinct types based on microscopic examination (Fig. 2A, B, C, D). - normal nuclei, pyknotic nuclei, and pale-staining/ poorly staining nuclei. Uninfected BMDMs from Optn^+/+^ and Optn^-/-^ mice showed similar gross cellular and nuclear morphology, and macrophages from both Optn^+/+^ and Optn^-/-^ mice showed aggregation and altered cellular morphology 8 h and 20 h post-infection. We observed a considerable reduction in the number of macrophages 8 h post-infection, with altered nuclear morphology and an increase in pyknotic and pale staining nuclei. Notably, Optn^-/-^ macrophages showed a significantly higher percentage of pale staining nuclei compared to Optn^+/+^ macrophages 20 h post-infection (Fig. 2E). We made similar observations with high MOI infection of BMDMs with *M. tb* H37Ra, and *M. bovis* BCG (data not shown). These findings suggest that mycobacterial infection alters the cellular and nuclear morphology of macrophages, which may have implications in the immune response and pathogenesis of mycobacterial infections.

**Figure 2:**
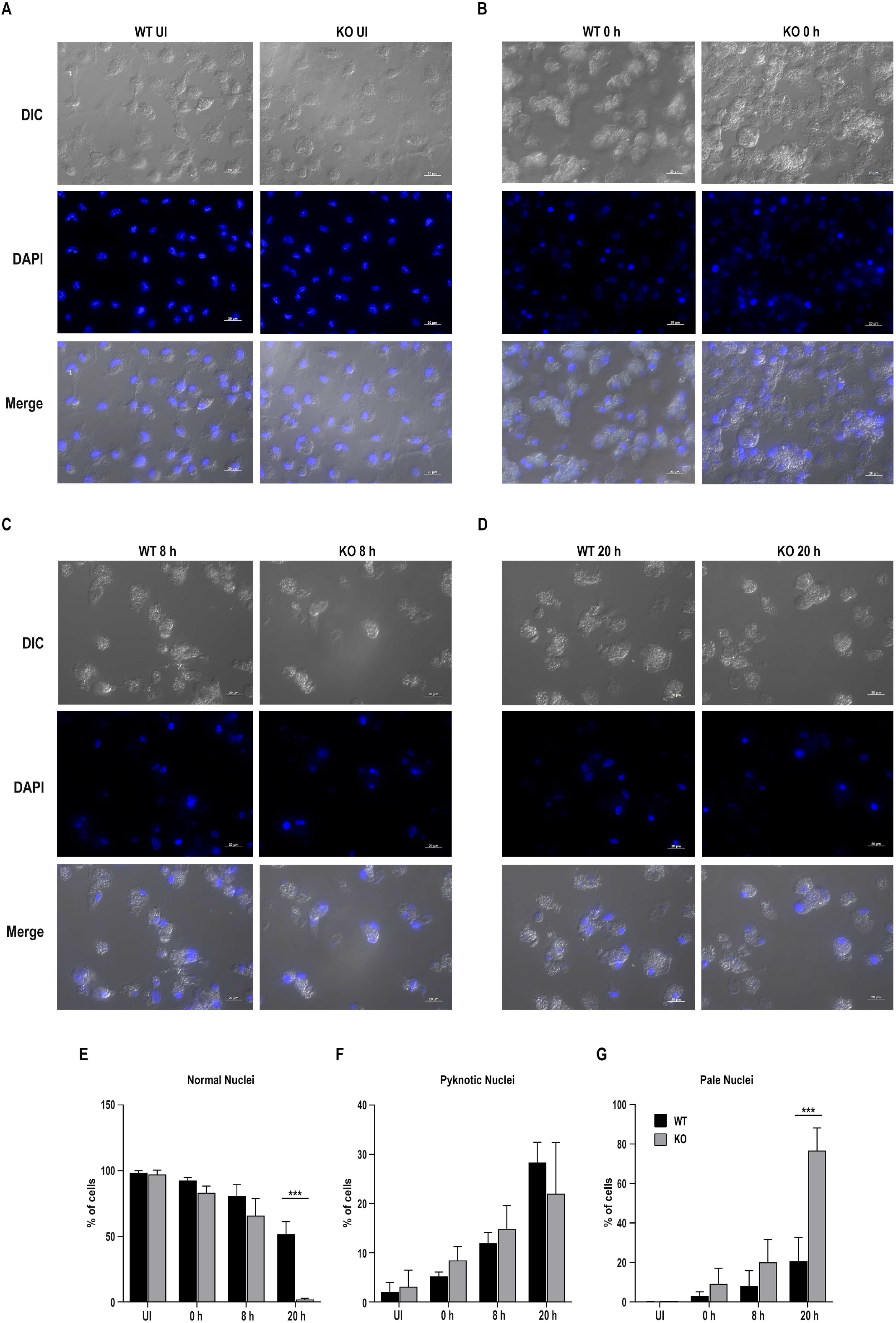
Microscopic observation of nuclear morphology changes in *M. smegmatis* infected BMDMs at 50:1 MOI. (A) Cellular and nuclear morphology of uninfected BMDMs from *Optn*^+/+^ and *Optn*^-/-^ mice. Cellular and nuclear morphology of *M. smegmatis* infected BMDMs from *Optn*^+/+^ and *Optn*^-/-^ mice, 0 h (B), 8 h (C), and 20 h (D) post infection. (E). Quantification of BMDMs based on nuclear morphology as into normal, pyknotic and pale staining nucleus. (n=4 independent experiments, minimum 300 cells per experiment per genotype assessed) Bars represent mean ± SD. ***p ≤ 0.001.

### Optineurin-deficient macrophages infected with *M. smegmatis* at High MOI induce diminished LC3 II levels

The observed changes in nuclear morphology suggested that upon mycobacterial infection, macrophages undergo time-dependent cellular changes that eventually lead to cell death. Since Optineurin is an autophagy receptor, we investigated changes in autophagy during mycobacterial infection. Induction of LC3II levels in macrophages from both Optn^+/+^ and Optn^-/-^ mice-derived BMDMs were observed post-infection with *M. smegmatis* (Fig. 3A), however, BMDMs derived from Optn^-/-^ mice showed reduced induction of LC3II 20 h post infection (Fig. 3B). This finding is consistent with the observation of increased cell death in Optn^-/-^ macrophages and suggests that altered autophagy may be one of the reasons for this cell death phenomenon. We also quantified the levels of activated caspase-3 in macrophages and found that the differences in these levels were not significant. This suggested that classical apoptosis did not play a significant role in the altered death of infected Optn^-/-^ macrophages.

**Figure 3:**
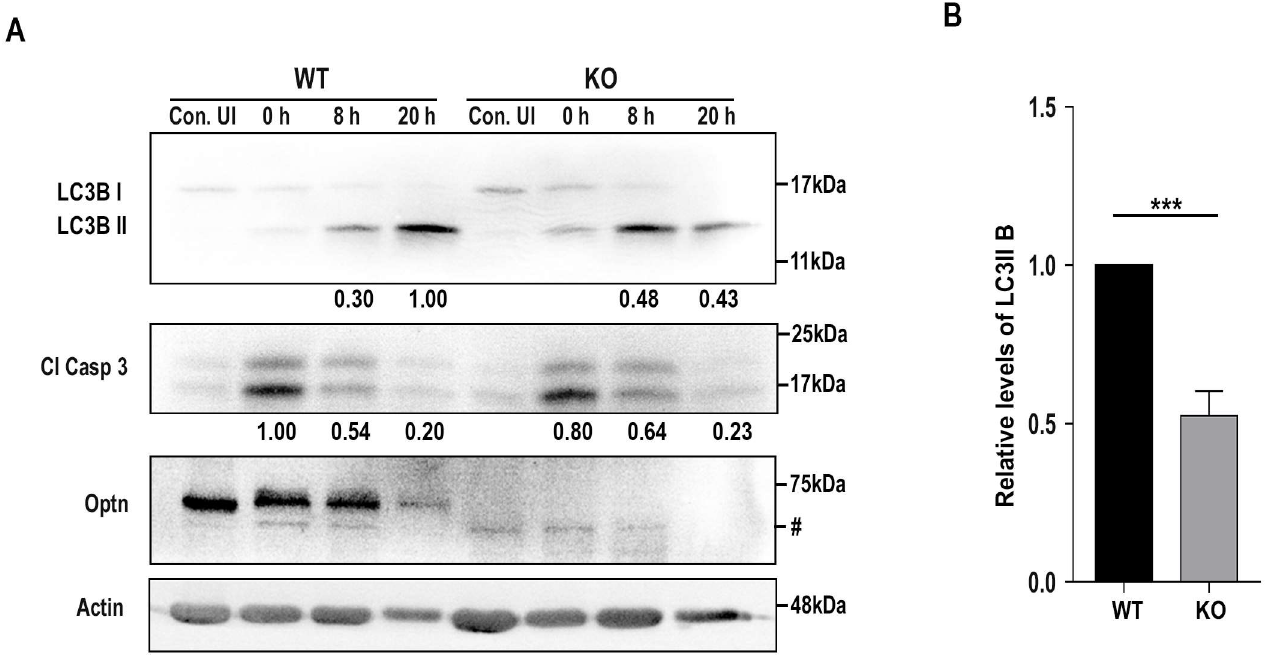
Assessment of autophagy in *M. smegmatis* infected Bone marrow-derived macrophages from *Optn*^+/+^ and *Optn*^-/-^ mice: (**A**) Representative western blot showing the levels of LC3, Cleaved Caspase 3, Optineurin, and β-Actin in *M. smegmatis* infected BMDMs isolated from *Optn*^+/+^ and Optn^-/-^ mice. (**B**) Bar diagram showing quantified levels of LC3 II, at 20 h post-infection. (n=4 independent experiments) Bars represent mean ± SD. ***p ≤ 0.001. ‘#’ indicates a non-specific band. Con. UI = Uninfected Control macrophages.

### Optineurin-deficient bone marrow-derived macrophages show altered inflammatory responses

Interleukin-1β (IL-1β) is a crucial cytokine produced and secreted by macrophages in response to infection. The protein is synthesized as a 32 kDa pro-protein and processed by Caspase-1 into an active 17.5 kDa form, which is then secreted into the bloodstream. IL-1β functions as a lymphocyte activating factor and a leukocytic pyrogen, mediating inflammatory responses. To investigate the impact of optineurin deficiency on the inflammatory response of macrophages, we treated primary BMDMs derived from Optn^+/+^ and Optn^-/-^ mice with 1μg/ml LPS derived from *E. coli* and measured the induction of Pro IL-1β. Western blot analysis showed that Optn^-/-^ macrophages had higher levels of Pro IL-1β compared to Optn^+/+^ macrophages after 24 h of LPS treatment (Fig. 4A). Statistical analysis revealed that the difference in Pro IL-1β levels was significant (Fig. 4B). These results suggest altered inflammatory response signalling in primary BMDMs under conditions of optineurin deficiency.

**Figure 4:**
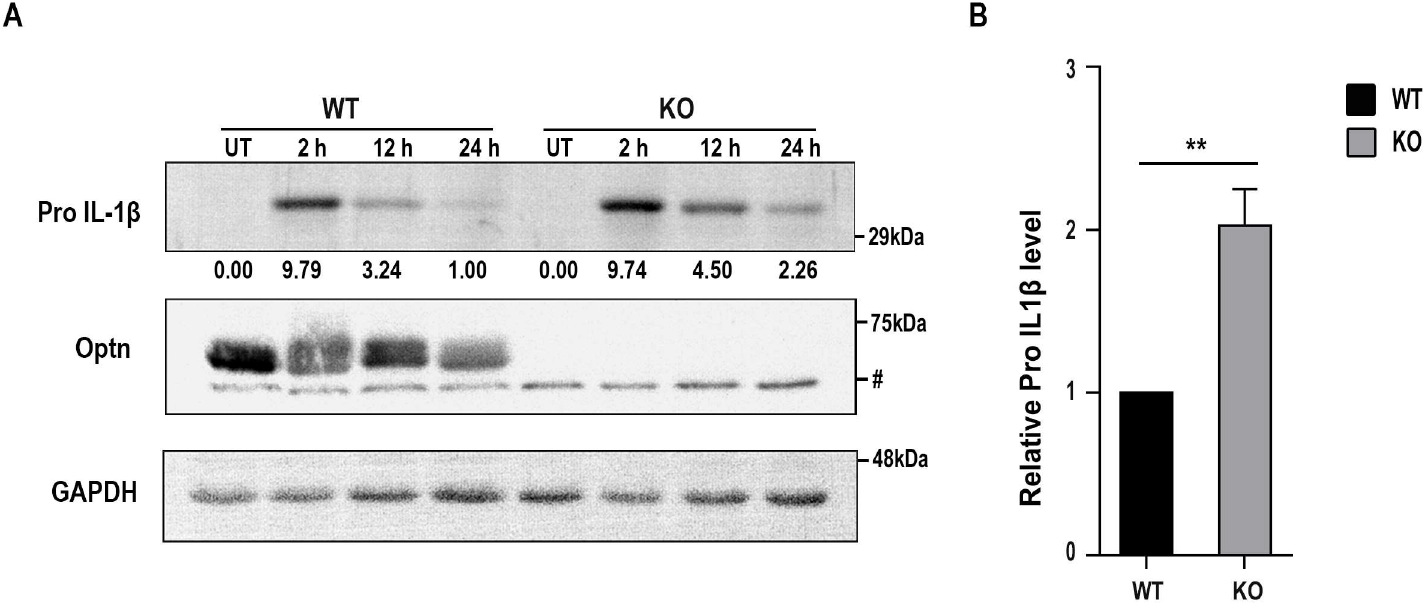
Levels of Pro-IL-1β in Bone marrow-derived macrophages from *Optn*^+/+^ and *Optn*^-/-^ mice: (**A**) Levels of Pro IL-1β in BMDMs isolated from *Optn*^+/+^ and Optn^-/-^ mice on treatment with 1μg/ml LPS. **B**) Bar diagram showing quantified levels of Pro IL-1β at 24h post LPS treatment (n=3 independent experiments). Bars represent mean ± SD. **p ≤ 0.01. ‘#’ indicates a non-specific band.

## DISCUSSION

A report investigating the role of optineurin in bacterial infection showed that optineurin restricts the growth of *Salmonella* in HeLa cells, consistent with its function as an autophagy receptor [7]. However, our results show that while optineurin deficiency did not alter CFU counts under low MOI *M. smegmatis* infection of BMDMs, under high MOI conditions, CFU counts in infected macrophages derived from *Optn*^-/-^ mice were lower than macrophages derived from *Optn*^+/+^ mice. It is probable that this results from the large increase in cell death in *Optn*^-/-^ mice-derived BMDMs, which then become freely permeable to the gentamycin in the culture medium, thereby killing intracellular *M. smegmatis*. As a consequence, bacteria in dead macrophages would not contribute to the CFU counts, leading to the lower CFU counts that we observe in infected *Optn*^-/-^ BMDMs.

Xenophagy clears pathogenic intracellular organisms using autophagic receptors such as Optineurin, to recognise the ubiquitinated bacteria that are destined to be degraded [36, 37]. The cargo destined for degradation is brought to LC3-positive phagophores called autophagosomes [38]. Optineurin appears to be important for pathogen clearance, as it was found to regulate the growth and replication of *Salmonella*, and Optineurin deficient mice were observed to be more susceptible to *Salmonella enterica* serovar *typhimurium* infection [7, 19]. The association between Optineurin and Mycobacterial infection has been a subject of recent investigations.

Optineurin expression was observed to be upregulated upon *M. marinum* infection in macrophages. In addition, a deficiency of Optineurin leads to increased susceptibility to *M. marinum* infection [39, 40]. Among various mycobacterial species *M. smegmatis* is recognized for its ability to induce the highest level of autophagy [41]. Since our focus was to investigate the role of the autophagy receptor optineurin during mycobacterial infection, selecting this as the test organism for our studies was the logical choice. Optineurin deficiency in BMDMs causes reduced macrophage autophagy, as observed by reduced LC3II levels upon *M. smegmatis* infection. It is known from the literature that a deficiency of optineurin leads to reduced autophagosome formation and reduces cargo selective autophagy [42, 43]. In our experiments, infection with *M. smegmatis* resulted in increased pale staining nuclei in optineurin deficient macrophages, which are likely to be dead macrophages. This cell death in infected macrophages coinciding with diminished autophagy, could be the cause of increased cell death in *Optn*^-/-^ mice derived BMDMs during *M. smegmatis* infection.

The observation that LPS treatment of optineurin-deficient macrophages leads to altered inflammatory stress response signalling, suggests that optineurin deficiency by itself predisposes cells to altered inflammatory signalling, and therefore leads to a poor infection response in BMDMs. Reduced expression of Optineurin is linked to impaired cytokine secretion by macrophages and altered inflammatory responses [44]. In addition, a reduction in Optineurin expression in mice led to diminished levels of pro-inflammatory TNFα in the serum, diminished cytokine secretion, diminished neutrophil recruitment to sites of acute inflammation, resulting in greater mortality after bacterial stimulation [45]. our findings show that Optineurin-deficient BMDMs are more sensitive to *M. smegmatis* induced cell death indicating that optineurin-mediated modulation of the infection response has a cytoprotective function against *M. smegmatis* induced cell death.

Exposure of BMDMs to high MOI *M. smegmatis* leads to rapid and substantial bacterial uptake. In contrast to low MOI infection this high bacterial load results in BMDMs become overwhelmed, leading to their death. Our results show that Optineurin deficient BMDMs have defective pro-inflammatory signalling. The excess of immune signalling molecules during high MOI infection can lead to cytotoxic effects on BMDMs themselves, contributing to increased cell death. BMDMs are known to generate reactive oxygen species as a defence mechanism against invading pathogens. OPTN is expressed in bone and neuronal cells and has been reported to protect cells from ROS-induced cell damage [46, 47]. However, in high MOI mycobacterial infection, the excessive production of ROS can cause cellular damage and contribute to death of BMDMs, with the absence of optineurin intensifying these effects.

Taken together, our results demonstrate that optineurin is critical for regulating macrophage responses during mycobacterial infection. Optineurin deficiency alters cytokine production and autophagy, key events in cellular responses to infection, highlighting both its cytoprotective function, and its potential therapeutic importance in the treatment of TB.

## CONFLICT OF INTEREST

The authors declare that there are no conflicts of interest.

## FUNDING INFORMATION

This work was supported by grant from the Council of Scientific and Industrial Research (MLP 0144)), Government of India (CSIR) (to T. R. R.), and by a J.C. Bose National Fellowship grant (SR/S2/JCB-41/2010) from the Department of Science and Technology, Government of India (to G.S.). The funders had no role in study design, data collection and analysis, decision to publish, or preparation of the manuscript. G. R. was supported by a fellowship from the Indian Council of Medical Research, C.V.Y. was supported by a Research Associateship from the the Department of Biotechnology, Government of India.

## AUTHOR CONTRIBUTIONS

T. R. R. and G.S. designed the study. G.R. and C.V.Y. performed the experiments. T. R. R., G.S., and G.R. analysed the data. T.R.R., G.S., and G. R. wrote the manuscript.

## Notes

### Competing Interest Statement

The authors have declared no competing interest.

